# Modified Pearson Correlation Coefficient for Two-color Imaging in Spherocylindrical Cells

**DOI:** 10.1101/300517

**Authors:** Sonisilpa Mohapatra, James C. Weisshaar

## Abstract

The revolution in fluorescence microscopy enables sub-diffraction-limit (“superresolution”) localization of hundreds or thousands of copies of two differently labeled proteins in the same live cell. In typical experiments, fluorescence from the entire three-dimensional (3D) cell body is projected along the *z*-axis of the microscope to form a 2D image at the camera plane. For imaging of two different species, here denoted “red” and “green”, a significant biological question is the extent to which the red and green spatial distributions are positively correlated, anti-correlated, or uncorrelated. A commonly used statistic for assessing the degree of linear correlation between two image matrices **R** and **G** is the Pearson Correlation Coefficient (PCC). PCC should vary from –1 (perfect anti-correlation) to 0 (no linear correlation) to +1 (perfect positive correlation). However, in the special case of spherocylindrical bacterial cells such as *E*. *coli* or *B*. *subtilis*, we show that the PCC fails both qualitatively and quantitatively. PCC returns the same +1 value for 2D projections of distributions that are either perfectly correlated in 3D or completely uncorrelated in 3D. The PCC also systematically underestimates the degree of anti-correlation between the projections of two perfectly anti-correlated 3D distributions. The problem is that the projection of a random spatial distribution within the 3D spherocylinder is non-random in 2D, whereas PCC compares every matrix element of **R** or **G** with the constant mean value *R* or *G*. We propose a modified Pearson Correlation Coefficient (MPCC) that corrects this problem for spherocylindrical cell geometry by using the proper reference matrix for comparison with **R** and **G**. Correct behavior of MPCC is confirmed for a variety of numerical simulations and on experimental distributions of HU and RNA polymerase in live *E*. *coli* cells. The MPCC concept should be generalizable to other cell shapes.

## INTRODUCTION

In widefield and superresolution fluorescence microscopy of eukaryotic and prokaryotic cells, the fluorescent species occupy a three-dimensional (3D) volume. In typical usage, the laser illuminates the entire thickness of the cell (“epi illumination”). The microscope then projects fluorescence from a 3D source along the *z* axis to form a two-dimensional (2D) image at the *xy* camera plane. For two-color imaging of two different species, herein called the “red species” and the “green species”, an important biological question is the degree to which the red and green spatial distributions are positively correlated, anti-correlated, or uncorrelated with each other. Positive correlation may suggest binding to a common cytoplasmic element such as a membrane or the chromosomal DNA or common sites of production, action, or degradation. Negative correlation may suggest a physical or biochemical mechanism that sequesters red and green species from each other (1, 2).

The Pearson correlation coefficient (PCC) (3, 4) is one of the most commonly used statistical tools to measure the degree of linear correlation between two data sets X and Y:

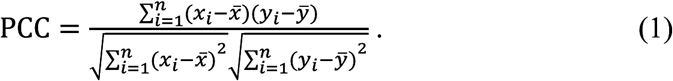

Here (*X_i_*, *y_i_*) are individual paired samples from the data sets X and Y and *n* is the total number of pairs; *x̄* and *ȳ* are the mean values of the samples in data sets X and Y. With the advent of two-color superresolution fluorescence microscopy, the PCC is increasingly used as a statistic for quantifying the degree of correlation between the subcellular distributions of two distinguishable species. For image matrices **R** (red channel) and **G** (green channel), the formula for PCC becomes:

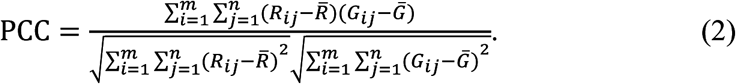

Here *m* and *n* are the number of rows and columns in the image matrices; there are *m* x *n* total pixels in each image. The *R_ij_* and *G_ij_* are the corresponding intensities of pixel *ij* in **R** and **G**; for superresolution images these are integers (counts/pixel). *R* and *G* are the mean pixel intensities of **R** and **G**. In the PCC formula, all elements of the reference matrix with which **R** or **G** is compared have the same value. The value *R* (or *G*) is subtracted from each individual pixel intensity *R_ij_* (or *G_jy_*), yielding both positive and negative difference intensities (*R_ij_* – *R*) and (*G_jy_* – *G*). Thus the product in the PCC numerator provides information about the correlation between deviations of *R_ij_* from *R* and deviations of *G*_*jy*_ from *G*. The denominator normalizes PCC so that it always lies in the range –1 to +1. Ideally, PCC = 1 indicates two perfectly linearly correlated images for which each red pixel *ij* deviates from the red mean in direct proportion to the deviation of the corresponding green pixel *ij* from the green mean. PCC = 0 indicates two linearly uncorrelated images. PCC = –1 indicates two perfectly anti-correlated images (red and green deviations of equal magnitude but of opposite sign). A PCC value significantly different from zero is a measure of the degree to which two distributions are correlated or anti-correlated as compared with the null hypothesis of PCC = 0, corresponding to two uncorrelated, random distributions.

The ImageJ software (5) extensively used for image analysis in the field of fluorescence microscopy provides Coloc2 and JaCoP plugins (6) that enable the user to calculate PCC between two images. In the recent literature, PCC has been used to characterize the correlation in 2D spatial distributions of two fluorescently labeled proteins in both bacterial cells (7-9) and eukaryotic cells (10-16). McDonald and co-workers recently catalogued some common pitfalls in the use of PCC on eukaryotic cells (12).

We are particularly interested in small, rod-shaped, approximately spherocylindrical bacterial cells such as *E*. *coli* and *B*. *subtilis*, whose typical length is *L_cell_* ∼ 4 μm and whose diameter is 2*r* ∼ 1 μm. For the most common shapes of bacteria (spherical, rod-shaped and spiral), the standard PCC procedure fails both qualitatively and quantitatively. Spherocylinders have strong curvature at the two endcaps and in the cylindrical region. As a result, the projection of molecules randomly distributed in a 3D spherocylindrical volume does not form a random distribution in 2D. In Fig. 1, we illustrate the 2D projection of 5000 molecules that are distributed randomly in a 3D spherocylinder with dimensions similar to that of an *E*. *coli* cell in good growth conditions. The endcap regions and the edges of the spherocylinder project a smaller volume onto the camera plane, and thus have fewer counts/pixel in the 2D image than the central cylindrical region. This effect is clear in the pixelated 2D localization density maps shown in Fig. 1*C-E*. Pixels in the 2D projection of a random 3D distribution vary in intensity by a factor of five or more, depending on the chosen pixel size. The variations are highly systematic.

**Figure 1.**
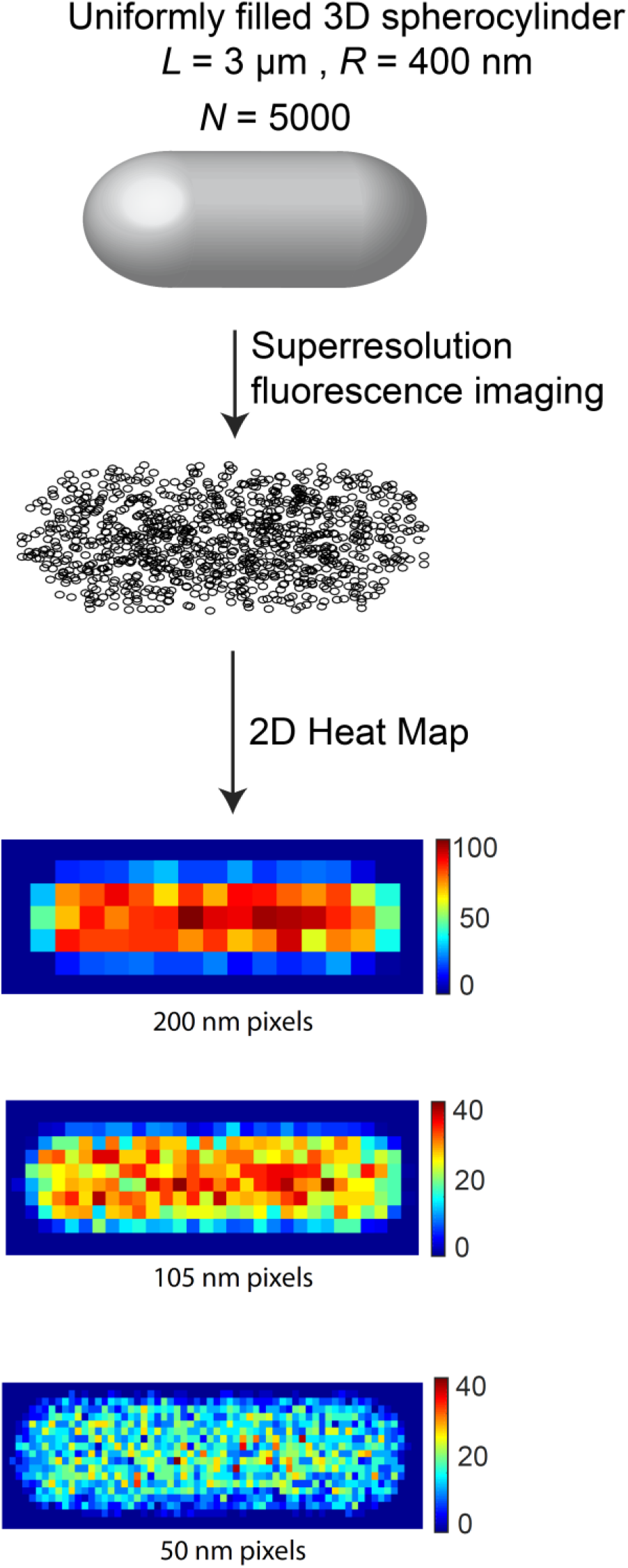
Schematic of method for obtaining a 2D pixelated image from 3D distribution of molecules within a spherocylinder. **(A)** Uniformly filled spherocylinder representing a bacterial cell cytoplasm. **(B)** 2D projection of 5000 molecules distributed randomly in the 3D spherocylinder obtained by superresolution fluorescence imaging. **(C)** – **(E)** 2D localization probability density heat maps of imaged molecules with individual pixel sizes of 200 nm,105 nm, and 50 nm.

Consequently, the PCC reference matrix used for comparison with **R** and **G** is inappropriate. The PCC difference intensities (*R_ij_* – *R*) and (*G_ij_* – *G*) for pixels at the edges and end caps are both systematically negative, *i*.*e*., strongly biased towards having fewer molecules/pixel than the mean value in a 2D projection of a 3D random distribution. In those regions, the products (*R_ij_* – *R*)(*G_ij_* – *G*) are systematically positive. Similarly, the difference intensities of the pixels in the central region of the spherocylinder are systematically positive, strongly biased towards having more molecules/pixel than the mean of a projection of a 3D random distribution. In that region, the products (*R_ij_* – *R*)(*G_ij_* – *G*) are again systematically positive. For two uncorrelated, random distributions in 3D, this causes the traditional PCC of the 2D projection to incorrectly approach +1, not the desired result of zero. The same systematic positive bias causes the traditional PCC to underestimate the degree of anti-correlation between two perfectly anti-correlated images, as we will show.

In the following sections, we describe a procedure for calculating what we call the modified Pearson correlation coefficient (MPCC) in the special case of interest, spherocylindrical bacterial cells like *E*. *coli* and *B*. *subtilis*. The procedure should prove useful for both widefield and superresolution images, and in principle it could be adapted to other cell shapes. We use numerical simulations to show that MPCC properly becomes zero for two uncorrelated, random distributions, approaches –1 for two perfectly anti-correlated images, and approaches +1 for two perfectly correlated images. We also provide guidance for pixelation of superresolution images and show how to determine the probability *p* that a measured non-zero MPCC did not arise from two uncorrelated, random 3D distributions. We conclude with an experimental example of a significantly positive MPCC between superresolution images of RNA polymerase and of the DNA-binding protein HU in live *E*. *coli*. The package of MATLAB codes required for calculating MPCC between two different molecules imaged in rod shaped cells such as *E*. *coli* and *B*. *subtilis* is available on GitHub: *https://github.com/SoniMohapatra/MPCC*.

## RESULTS

### The modified Pearson correlation coefficient MPCC

The MPCC of two images **R** and **G** is evaluated as follows:

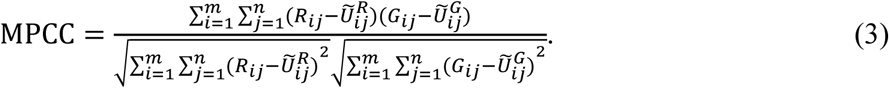

Here we have replaced *R* and *G* in Eq. 2 with the modified reference matrices
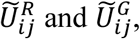
respectively.
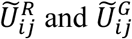
denote the intensity of pixel *ij* in the 2D projection of a large set of molecules distributed randomly in a 3D spherocylinder. The total number of molecules in **U͠**^*R*^ and **U͠**^*G*^ has been scaled to be the same as the total number of molecules in **R** and **G**, respectively.

In favorable conditions, superresolution imaging provides (*x*,*y*) spatial localization of hundreds or thousands of molecules per cell with spatial resolution of *σ*_*x*,*y*_ ∼ 20–50 nm. Conversion of these single molecule locations into 2D probability density maps requires selection of a pixel size; several examples are shown in Fig. 1*C*-*E*. The intensity in each pixel equals the total number of molecules assigned to it. The dependence of the calculated MPCC on the chosen pixel size and the number of imaged molecules is described later. These pixelated 2D maps for the red and green channels are denoted by **R** and **G**, the image matrices in Eq. 3.

To form the numerator of Eq. 3, we then subtract **U͠^R^** and **U͠^G^** from the corresponding image matrix in the red and green channels (**R** and **G**, respectively) to obtain the (unnormalized) difference matrices **Δ^R^** and **Δ^G^**. The resultant difference matrices have pixels with positive and negative values. Finally, to constrain MPCC to lie in the range +1 to −1, we normalize **Δ^R^** and **Δ^G^** so that the sum of the squares of individual pixel values in the difference matrix is 1. The resultant normalized 2D difference matrices are called **Δ̂^R^** and **Δ̂^G^** respectively. MPCC is obtained by taking the Frobenius inner product of the two normalized matrices **Δ̂^R^** and **Δ̂^G^** (Eq. 6 in Methods). A detailed step-by-step description of the methodology for obtaining MPCC is presented in the Methods section.

The MPCC ranges from +1 to −1, as does standard PCC. The MPCC for two images is +1 when the normalized difference matrices are perfectly linearly related, *i*.*e*., when
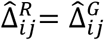
for every pixel *ij*. As a result, MPCC =
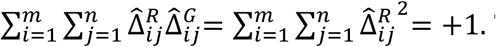
The MPCC is −1 when the normalized difference matrices are perfectly inversely related to each other, *i*.*e*.,
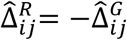
for every pixel. As a result,

MPCC =
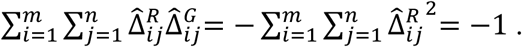
When the normalized difference matrices of two images are uncorrelated with each other, the MPCC is 0.

Next we carry out numerical simulations comparing MPCC with PCC for 2D projections from 3D spherocylinders in three cases: perfect 3D correlation that projects into perfect 2D correlation, perfect 3D anti-correlation that projects into perfect 2D anti-correlation, and uncorrelated, random 3D distributions. For all these examples, the **R** and **G** image matrices have 10,000 molecules each. The spherocylinder has tip-to-tip length *L_cell_* = 3.5 μm and diameter 2*r* = 0.82 μm. The 2D pixel size in the image matrices **R** and **G** is chosen to be 200 nm in both dimensions, so that 75 pixels cover the 2D projection.

#### Perfect anti-correlation in 3D

To examine the case of two perfectly anti-correlated images, we have simulated 3D random distributions of 20,000 molecules confined to the spherocylindrical volume. The ∼10,000 molecules located in the left half of the spherocylinder are designated red; the ∼10,000 molecules located in the right half are designated green. This ensures that there is no spatial overlap of molecules in the red and green channels. We call this anti-correlation Case I. For such strong spatial anti-correlation, we should expect MPCC = −1. An example of the corresponding 2D image matrices **R** and **G** is shown in Fig. 2*A*. In Fig. 2*B*, *C*, we have compared the reference matrices and the key normalized difference matrices the products of whose corresponding elements enter the traditional PCC (Eq. 2) and the new MPCC (Eq. 3).

**Figure 2.**
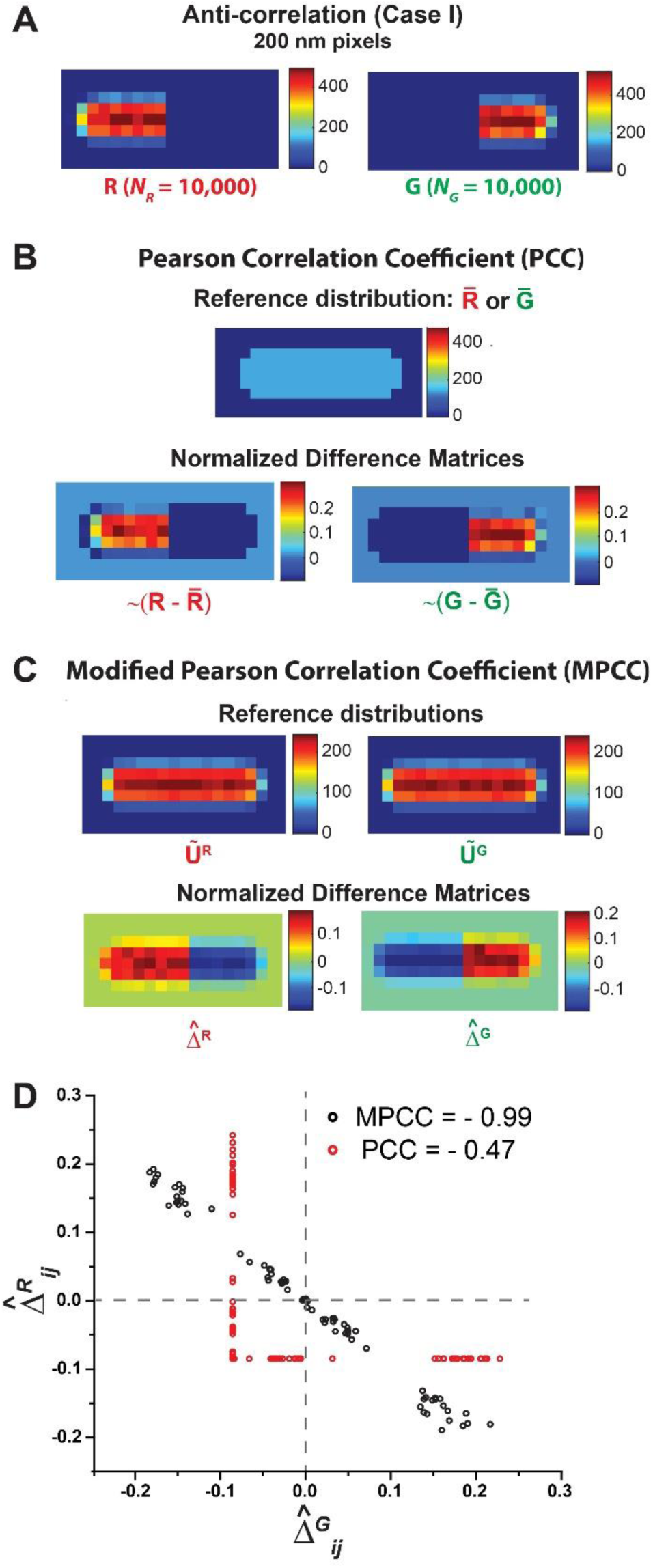
Scheme for calculating PCC and MPCC for two representative images **R** and **G** that are perfectly anti-correlated in both 3D and 2D. **(A)** Heat maps of **R** and **G** with 200 nm pixels. Each image comprises ∼10,000 molecules. Color scale indicates the number of molecules in each pixel. **(B)** Standard PCC calculation. *Top*: The 2D uniform reference distribution **R** or **G** that is subtracted from images **R** or **G**. *Bottom*: Normalized difference matrices ∼(**R – R**) and ∼(**G – G**) obtained after subtraction. The Frobenius inner product of these two difference matrices gives the PCC. **(C)** Modified PCC calculation. *Top*: Reference distribution **U͠^R^** and **U͠^G^**, which are 2D projections of 3D random distributions of 100,000 molecules within the spherocylinder and normalized to have a total of 10,000 molecules. These are subtracted from images **R** and **G**, respectively. *Bottom*: Normalized difference matrices **Δ̂^R^**and **Δ̂^G^** obtained after subtraction. The Frobenius inner product of these two difference matrices gives the MPCC. **(D)** Scatter plot of individual normalized difference matrix elements for PCC (*Red*) and for MPCC (*Black*). The MPCC elements are negatively correlated within the noise level, while the PCC elements are not. The resulting MPCC and PCC values are –0.99 and –0.47, respectively.

For the traditional PCC (Fig. 2*B*), there are ∼10,000 molecules of each color distributed in a cell area covering 75 pixels. As in Eq. 2, we subtract the mean pixel intensity *R* = 133.3 and *G* = 133.3 from each individual pixel intensities *R_ij_* and *G_ij_*. The resulting normalized difference matrices,
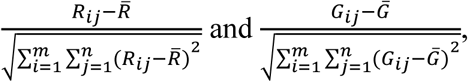
are depicted as heat maps labeled ∼(**R –** **R**) and ∼(**G – G**) in Fig. 2*B*. These are the PCC analogues of
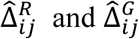
in the MPCC equation. In the left half of the spherocylinder, the red difference matrix has a thin shell of systematically negative values (endcap and edge pixels) and a central core of systematically positive values. When multiplied by the corresponding elements of the left half of the green difference matrix, which contains all equal negative elements, the contributions to PCC will be positive and negative, respectively. The same type of systematically positive and negative contributions will arise from the right half of the spherocylinder. The resulting red and green contributions to PCC are not linearly anti-correlated. This is seen clearly in Fig. 2*D*, where we show a scatter plot of the individual red normalized differences vs the corresponding green normalized differences. The net result is PCC = –0.47, suggesting only partial anti-correlation of the two spatial distributions even though they are completely anti-correlated in both 3D and 2D.

In contrast, the MPCC formula of Eq. 3 subtracts from each pixel the proper 2D contribution of the projection of a smooth 3D random distribution (Fig 2*C*). The resulting normalized difference matrices **Δ̂^R^** and **Δ̂^G^** are also depicted in Fig. 2*C*. The scatter plot of individual difference matrix elements
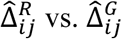
in Fig. 2*D* shows the expected strong linear anti-correlation for all pixels. The resulting MPCC is −0.99, very close to the expected value of –1.

In SI Text S1 in the Supplemental Information, we examine two additional examples of perfect anti-correlation. In anti-correlation Case II shown in Fig. S1 in the Supplemental Information, the two endcap regions are occupied by ∼10,000 red molecules and the central region is occupied by ∼10,000 green molecules. Again, the normalized difference matrix elements are linearly anti-correlated and the calculated MPCC is –0.99. In anti-correlation Case III (Fig. S1), the ∼10,000 red molecules occupy the leftmost 2/3 of the spherocylinder volume while the ∼10,000 green molecules occupy the rightmost 1/3. The result is the same. The advantages of MPCC vs traditional PCC are apparent.

#### Perfect Positive Correlation in 3D and 2D

When the red and green 3D spatial distributions are perfectly positively correlated, so will be their 2D projections. As described before, an MPCC value of +1 is expected for a case of perfect correlation in the 2D projections. The same is true of the traditional PCC. To examine the case of two perfectly correlated images, we have simulated 3D random distributions of 20,000 molecules confined to the spherocylindrical volume. The ∼10,000 molecules located in the left half of the spherocylinder are designated red; the molecules in the right half are deleted. We then independently simulated another 20,000 molecules distributed randomly in a 3D spherocylinder. The ∼10,000 molecules located in the left half of the spherocylinder are designated green; the molecules in the right half are again deleted. The resulting 3D distributions are projected into 2D and pixelated to yield the image matrices depicted in Fig. S2*A* in the Supplemental Information. We calculate the MPCC = +0.99 between these two distributions, very close to the anticipated value of 1. The resulting normalized difference matrices **Δ̂^R^** and **Δ̂^G^** obtained during evaluation of MPCC are depicted in Fig. S2*C*. The scatter plot of individual matrix elements
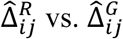
in Fig. S2*D* shows the expected strong linear correlation for all pixels. Similarly, the scatter plot of individual normalized difference matrix elements analogous to
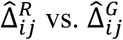
for PCC in Fig. S2*D* shows the expected strong linear correlation for all pixels. If *R_ij_* = *G_ij_* and *R* = *G*, then PCC = 1. Therefore, for two fluorescence images that are perfectly correlated in 3D, both the MPCC and the PCC will be +1 within the statistical noise.

#### Random Distributions in 3D

Two independent, uncorrelated, random distributions should have a Pearson correlation coefficient of 0 within the statistical noise. In the numerical tests, we have randomly distributed 10,000 red molecules and 10,000 green molecules in 3D within the spherocylinder. The two random distributions are generated independently, so we expect them to be uncorrelated with each other. We add appropriate localization errors *σ_R_* = 50 nm and *σ_G_* = 50 nm and then project the “measured” positions into the xy-plane. PCC and MPCC between the two 2D projection matrices (Fig. 3*A*) will be compared.

**Figure 3.**
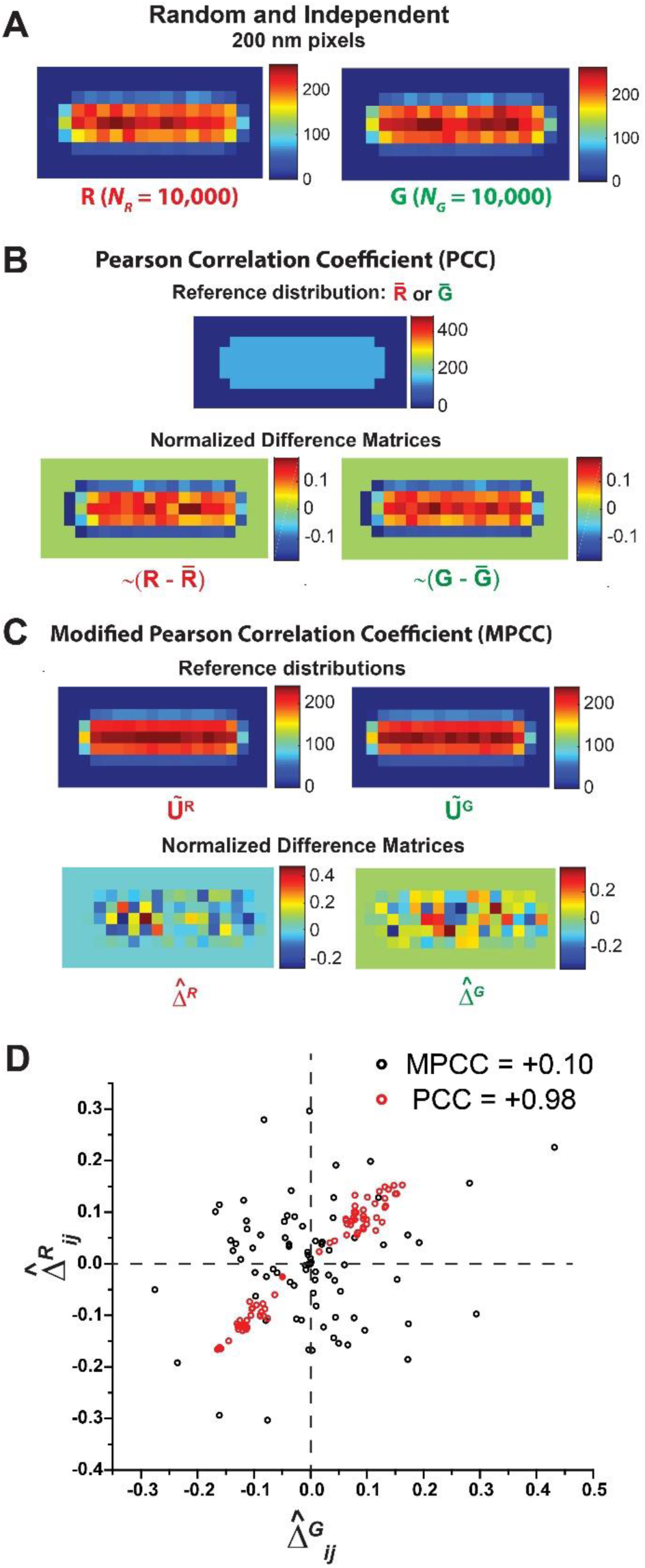
Scheme for calculating PCC and MPCC for two representative projected images **R** and **G** arising from two random and independent distributions in 3D. **(A)** Heat maps of **R** and **G** with 200 nm pixels. Each image comprises ∼10,000 molecules. Color scale indicates the number of molecules in each pixel. **(B)** Standard PCC calculation. *Top*: The 2D uniform reference distribution **R** or **G** that is subtracted from images R or G. *Bottom*: Normalized difference matrices ∼(**R – R**) and ∼(**G – G**) obtained after subtraction. **(C)** Modified PCC calculation. *Top*: Reference distribution **U͠^R^** and **U͠^G^**, which are 2D projections of3D random distributions of 100,000 molecules within the spherocylinder and normalized to have a total of 10,000 molecules. These are subtracted from images **R** and **G**, respectively. *Bottom*: Normalized difference matrices **Δ̂^R^**and **Δ̂^G^** obtained after subtraction. **(D)** Scatter plot of individual normalized difference matrix elements for PCC (*Red*) and for MPCC (*Black*). The MPCC elements are randomly distributed, while the PCC elements are positively correlated. The resulting MPCC and PCC values are +0.10 and +0.98, respectively.

The resulting reference matrices and normalized difference matrices for PCC and for MPCC are depicted in Fig. 3*B and C* respectively. The scatter plots of
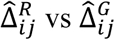
for MPCC and of their analogues for PCC are shown in Fig. 3*D*. The data indeed appear uncorrelated for MPCC, but they are strongly positively correlated for PCC. The resulting calculated coefficients are MPCC = +0.10 and PCC = +0.98. The cause of the large, positive PCC value between two random 3D distributions was described in the Introduction. The 2D projections have matching regions of systematically positive and systematically negative deviations from the 2D mean values.

Finally, we tested whether the distribution of calculated MPCC outcomes for two independent random distributions is appropriately centered at zero and unbiased towards positive or negative values. For 200 trials, we calculated MPCC values between two 2D projections of 3D independent, random distributions of 10,000 red and 10,000 green molecules using the same 200 nm pixel size. We fit the resulting distribution (Fig. S3 in the Supplemental Information) to a Gaussian function. The mean of the best-fit Gaussian distribution is <MPCC> = +0.0041 and the standard error is σ_MPCC_ = 0.13. The mean is close to zero and the distribution is symmetric about zero, as hoped for. The probability that a particular trial would yield an MPCC of magnitude 0.10 or larger on either side of the Gaussian distribution is *p* = 0.44. The “measured” example MPCC of +0.10 (Fig. 3*D*) lies within 1σ of the mean; it was not a particularly unusual event.

#### Dependence of MPCC and its uncertainty on pixel size and total number of imaged molecules

Before evaluating MPCC between two superresolution images, the pixel size in the 2D localization density maps must be chosen. For a fixed cell size, the smaller the pixel size, the greater will be the total number of pixels *N_p_*. We have shown in SI (Fig. S4 in the Supplemental Information) that for a fixed number of localizations *N_R_* = *N_G_* = 10,000 distributed randomly in 3D, as the pixel size decreases (and *N_p_* increases) the width of the distribution of MPCC values becomes narrower. All the MPCC distributions for uncorrelated images are symmetric and centered about 0 and well fit by a Gaussian function. For these random, uncorrelated 3D distributions, the standard deviation of the Gaussian MPCC distributions scales as
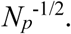
This scaling holds even for *N_R_* and *N_G_* as small as 500.

Narrower widths of the MPCC distribution from random 3D distributions generally provide greater statistical confidence that a non-zero measured value of MPCC is significantly different from zero. This argues for fine pixelation. In practice, we suggest simulating the distribution of MPCC values between the 2D projections of 3D random distributions using the same number of molecules as were imaged in the red and green channels and the same pixel size chosen for **R** and **G**. This enables assignment of a probability *p* that the measured MPCC arose from two random 3D distributions. If *p* is unacceptably large, finer pixelation of both experimental and simulated locations may decrease *p*.

However, for non-random 3D distributions such as the completely anti-correlated distribution of Fig. 2 or the positively correlated distribution of Fig. S2, it is important not to pixelate so finely that the matrices **R** and **G** become too sparse. In the case of anti-correlated **R** and **G**, this leads to false positive linear correlations between
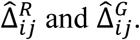
The zeroes appearing in the left-hand region of **R** positively correlate with the empty regions of **G**. Similarly, the zeroes arising due to sparseness in the right-hand region of **G** positively correlate with the empty regions of **R**. These systematically bias the MPCC for truly anti-correlated distributions towards more positive values, underestimating the degree of linear anti-correlation. We explore this effect numerically in Fig. S5 in the Supplemental Information. For a given pixel size, the mean MPCC moves closer to the expected value of −1 for two anti-correlated images as the number of imaged molecules increases. In practice, we suggest carrying out numerical simulations of perfectly anti-correlated images using values of *N_R_* and *N_G_* that match experiment. The pixel size chosen for analysis of the experimental data should be the smallest pixel size for which the mean MPCC for perfectly anti-correlated images is acceptably close to −1. In the numerical example of Fig. 2, with 10,000 molecules distributed over 75 pixels, the mean occupancy was 133 molecules/pixel, which yielded MPCC = –0.99. As a rule of thumb, it appears that if the mean occupancy is ∼7 copies/pixel (∼14 copies per pixel in the occupied halves of the case in Fig. 2), then the MPCC will be about –0.9.

For similar reasons, for two positively correlated images we expect that MPCC will systematically underestimate the degree of positive correlation as the red and green matrices become sparse. In the case of positively correlated **R** and **G** (Fig. S2), the zeroes appearing in the images due to sparseness are not positively correlated. The sparseness in number of molecules due to finer pixelation leads to false negative linear correlations between
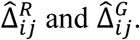
This leads to systematic negative deviations of the calculated MPCC from the expected value of +1. We investigated the mean occupancy/pixel that is required for the calculated MPCC between strongly positively correlated images to be ∼0.9, close to the expected value of +1. As shown in Fig. S5, a mean occupancy of ∼7 copies/pixel yields MPCC values of about +0.9.

Based on our investigations on anti-correlated and positively correlated images described above, we estimate that the minimum mean occupancy should be ∼7 copies/pixel for MPCC to give reasonably accurate results. The pixel size should be chosen keeping the minimum mean occupancy/pixel in mind.

### Experimental example of MPCC from superresolution images of RNAP and HU in *E*. *coli*

To test our MPCC concept on real experimental data, we performed two-color superresolution fluorescence imaging of RNA polymerase and HU in live *E*. *coli* cells. RNAP is primarily located in the nucleoid region because of its frequent specific and non-specific interactions with chromosomal DNA (17). HU is a DNA binding protein that should also localize within the nucleoids (18, 19). We expect significant positive correlation between the spatial distributions of RNAP and HU and therefore a positive value of MPCC.

For superresolution co-imaging of RNAP and HU in live *E*. *coli* cells, we constructed a strain where the gene coding for the fluorescent protein YFP (observed in the green channel) (20) is fused to the C terminus of the endogenous *rpoC* gene in *E*. *coli* VH1000. Single copies are imaged using the reversible photobleaching method described earlier (21). An inducible plasmid that expresses HU labeled with the photoactivatable fluorescent protein PAmcherry (22) (observed in the red channel) was introduced into the same strain. The cells were grown in EZ rich defined medium at 30°C, plated on a glass coverslip, and imaged with 30 ms exposure time. The details of strain construction, growth conditions, and imaging conditions are described in SI Text S3 in the Supplemental Information.

To obtain a useful number of imaged copies, we combine locations of red HU and green RNAP molecules from different cells of essentially the same length. The imaged cells were sorted by tip-to-tip length based on phase contrast images in order to avoid broadening of spatial distribution of molecules due to the range of cell lengths. For the analysis, we chose cells of length 3.6 to 3.8 μm, the bin with the highest number of imaged cells. The resulting composite distribution of spatial localizations of *N_G_* = 6570 RNAP-YFP and *N_R_* = 8436 HU–PAmcherry molecules from 11 cells pixelated to 105 nm (279 total pixels) is illustrated in Fig. 4*A*. The mean number of molecules per pixel is ∼25 and ∼30 for the RNAP and HU channels respectively. The corresponding 1D projected axial distributions are compared in Fig. 4*B*. The raw data indeed suggest significant positive correlation between the two distributions.

**Figure 4.**
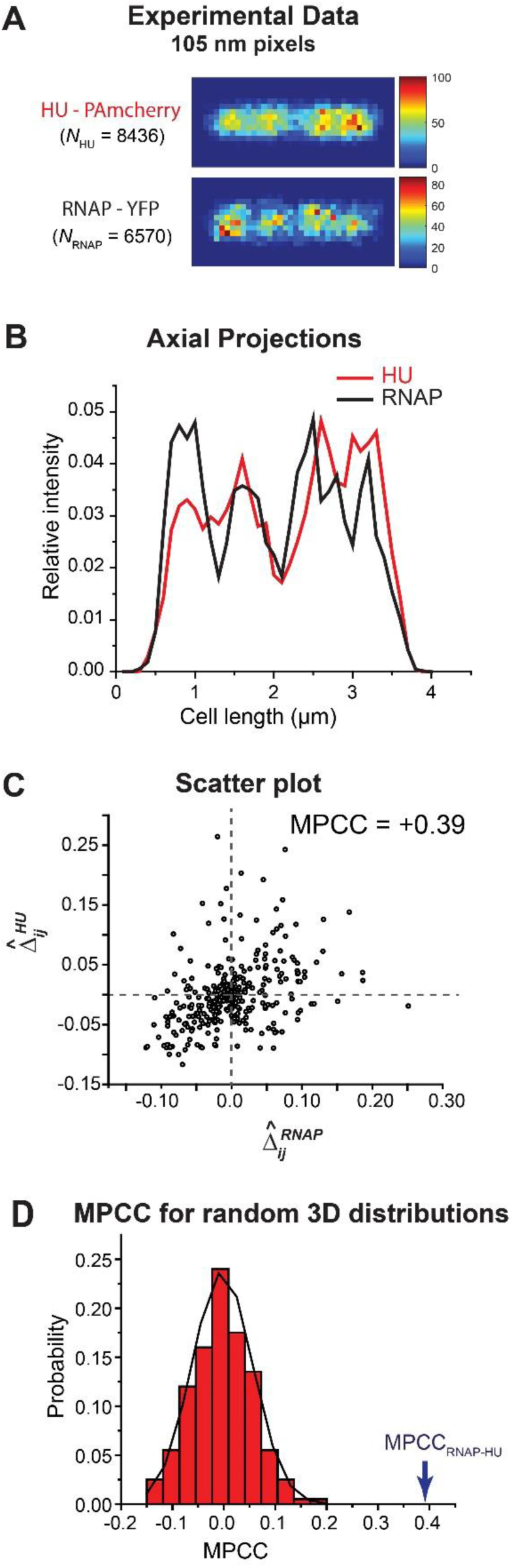
**(A)** Experimental 2D localization probability density maps of 8436 HU–PAmcherry molecules (*Top*) and 6570 RNAP–YFP molecules (*Bottom*). Composite of data from 11 cells of tip-to-tip length *L_cell_* in the range 3.6 to 3.8 μm. The color scale indicates the number of molecules in each pixel. **(B)** Axial probability density distributions of the imaged molecules. **(C)** Scatter plot of individual normalized difference matrix elements for MPCC, 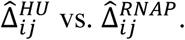
Plot shows significant visual evidence of positive correlation; the calculated MPCC is +0.39. **(D)** Histogram of 200 MPCC values calculated for pairs of independent, random 3D distributions using the same number of HU and RNAP copies and the same pixelation as the experimental data. Best fit to a Gaussian curve has <MPCC> = –0.0030 and σ = 0.061 (*Black* curve). The experimental MPCC (arrow) lies at +6.4σ, making it highly improbable that two random distributions would produce such a large, positive result.

For evaluation of MPCC we simulated two random distributions of 100,000 molecules each, corresponding to the RNAP (green) and HU (red) channels, using a spherocylinder whose dimensions match those of the chosen cells. The resulting reference images are normalized to have same number of molecules as imaged RNAP and HU. For accurate estimation of the cytoplasmic radius *r* of the imaged cells in the chosen length bin, we also imaged photoactivable Kaede molecules (23, 24), believed to distribute homogenously in the cytoplasmic volume (25). The detailed procedure is described in SI Text S4 in the Supplemental Information. The resulting cell length is *L_cell_* = 3.74 μm; the diameter is 2*r* = 0.82 μm (Fig. S6 in the Supplemental Information). The two simulated 3D random distributions incorporated localization errors σ_RNAP_ = 38 nm and σ_HU_ = 60 nm, determined by the intercepts of MSD plots (Fig. S7 in the Supplemental Information). We followed the procedure described above to calculate MPCC = +0.39. The scatter plot of
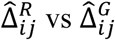
(Fig. 4*C*) also indicates significant positive correlation.

The final step estimates the probability *p* that a value of MPCC = +0.39 or larger would be obtained from two random 3D distributions with the same number of imaged molecules and the same pixel size used for the experimental data. In Fig. 4*D*, we show a histogram of the outcomes of 200 such simulations. The best-fit Gaussian distribution has a mean value <MPCC> = −0.0030 and standard error σ_MPCC_ = 0.061. The measured MPCC value lies 6.4σ_MPCC_ away from zero. Under the assumption that the statistics of the simulated MPCC trials are Gaussian, the probability that two random 3D distributions would produce an MPCC value of magnitude 0.39 or larger on either side of the Gaussian curve is *p* ∼ 1.6 × 10^−10^. Thus we can reject the null hypothesis that MPCC = +0.39 arose from two random, uncorrelated 3D distributions and assert significant positive correlation between the RNAP and HU distributions with very high confidence.

The choice of pixel size does affect the calculated MPCC. For 200 nm pixels (*N_p_* = 77 total pixels), the experimental MPCC is +0.46. The corresponding simulations of two of two random distributions gave <MPCC> = 0.0082 and σ_MPCC_ = 0.12. In this case, the probability that two 3D random distributions would produce an MPCC value of magnitude 0.46 or higher on either side of mean of Gaussian curve is *p* ∼ 1.3 × 10^−4^ For 50 nm pixels (*N_p_* = 1178 total pixels), the experimental MPCC is +0.25. The corresponding simulations of two random distributions gave <MPCC> = 0.0027 and σ_MPCC_ = 0.033. In this case, the probability that two 3D random distributions would produce an MPCC value of magnitude 0.25 or higher on either side of mean of Gaussian curve is *p* ∼ 3.6 × 10^−14^. The estimated experimental MPCC decreases systematically as *N_p_* increases and the same data set is pixelated more finely, but the simulated σ_MPCC_ decreases more rapidly. As described previously, finer pixelation of both experimental and simulated locations helps to decrease *p* and reject the null hypothesis with greater confidence. In all three cases, the calculated MPCC lies many standard deviations away from zero. Thus we deem the conclusion of significant positive correlation between the RNAP and HU experimental distributions to be robust.

As suggested by the projected axial distribution of RNAP and HU (Fig. 4*B*), the two species are not completely correlated in space. There are several factors that may explain why the MPCC is significantly smaller than 1. We have averaged the data over 11 cells whose nucleoids have irregular shapes in 3D that are not axially symmetric and that vary from cell to cell. The limited number of molecules lends statistical fluctuations to each distribution. In addition, while RNAP and HU both bind to the DNA, they have different biological functions and should not be expected to have spatial distributions that correlate perfectly.

## DISCUSSION

The Pearson correlation coefficient is one of the statistics commonly used for quantifying the degree of linear correlation in pixel-by-pixel intensity between two different images (3, 26-28). Owing to simplicity of usage and availability in most image analysis software packages (ImageJ, Colocalizer Pro), PCC is used increasingly in the literature of two-color fluorescence microscopy. Because it is pixel-based, PCC can be applied to both widefield and superresolution images. For sub-pixel resolution images, point-location methods such as Ripley’s K (29-31) or pair correlation methods (32, 33) provide a global overview of the spatial distribution of points. These methods allow determination of whether the proteins are dispersed, clustered, or randomly distributed within the region of interest. All these methods have limitations for 2D projections of 3D locations and for small bacterial cells whose boundaries distort the meaning of random distributions. Moerner and co-workers have recently applied Ripley’s K to the case of HU proteins in *C*. *crescentus* and corrected the reference random distribution by methods similar to those we employ here (34).

For two-color, three-dimensional fluorescence microscopy (35, 36), the PCC can provide an accurate measure of linear correlation, assuming the 3D image matrices are sufficiently populated. However, by far the more common case of two-color microscopy projects the 3D spatial distributions onto the 2D camera plane.

The central point of this work is simple. For most cell shapes, random 3D spatial distributions (no spatial correlations) do not make random 2D projections. In the particular case of spherocylindrical cells, projections of random 3D distributions are skewed to have more molecules/pixel in the central region compared to the edges and the endcap regions (Fig. 1). This renders the standard PCC reference matrices (Eq. 2), whose elements are the constant values *R* and *G*, highly inappropriate. As a result, the standard PCC fails both qualitatively and quantitatively to describe the nature and degree of the spatial correlation. A calculated PCC value of +1 could equally well arise from perfectly correlated 3D images (Fig. S2) or from completely random 3D images (Fig. 3). For strongly anti-correlated images, the degree of anti-correlation will be systematically underestimated (Fig. 2).

In the special case of spherocylindrical cells, we have described a method for calculating a modified Pearson correlation coefficient (MPCC) that uses the 2D projection matrices
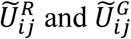
derived from independent 3D random distributions as the reference matrices with which the 2D image matrices **R** and **G** are compared (Eq. 3). The resulting MPCC is normalized to lie in the range –1 to +1. Within noise limitations, MPCC approaches 0 for the projections of two distributions that are independent and random in 3D, approaches –1 for two distributions that are perfectly anti-correlated in both 3D and 2D, and approaches +1 for two distributions that are perfectly positively correlated in both 3D and 2D. Additionally, we have used the new procedure to calculate a positive MPCC = +0.39 between experimentally obtained spatial localizations of individual RNAP and HU molecules in live *E*. *coli* cells (Fig. 4). Both RNAP and HU bind the chromosomal DNA, which occupies a subset of the cytoplasmic volume called the nucleoid. As expected, we obtain positive correlation that is significantly outside the range of model MPCC values computed for two uncorrelated distributions using the same pixel and copy number parameters as the experimental data.

While MPCC corrects a significant flaw in the standard PCC, it is important to note that for two images that are correlated or anti-correlated in 3D (*i*.*e*., not random), the MPCC applied to the 2D projections will typically underestimate the degree of correlation or anti-correlation in 3D. Projection from 3D to 2D always involves a loss of information. If the two 3D distributions are correlated or anti-correlated, their 2D projections will typically be less so. Our model correlated images (Fig. S2) and anti-correlated images (Figs. 2 and S1) are special cases in that they preserve perfect correlation or anti-correlation when projected from 3D to 2D. More irregular, less symmetric 3D distributions generally will not. This means that a 2D MPCC value that is not significantly different from zero does not imply the absence of 3D spatial correlations.

We have also shown how a small average number of molecules per pixel leads to undesirable zeroes in the **R** and **G** matrices, causing systematic errors in MPCC values (Fig. S5). For both truly anti-correlated images and truly correlated images, this effect diminishes the magnitude of MPCC (biasing it towards zero). The MPCC user needs to measure sufficient numbers of localizations and to make a knowledgeable choice of pixel size. As a first estimate, we prescribe a minimum of 7 molecules per pixels to obtain a trustworthy MPCC.

In earlier work applying PCC to eukaryotic cells, Dunn, *et*. *al* (12) warned against inclusion of empty extracellular regions in the image matrices **R** and **G**. Such extra zeroes alter the mean value in the references matrices and also artificially inflate the calculated PCC due to positive correlations between the empty regions in both matrices. They suggested carefully outlining only the regions of space that are occupied by the cells of interest. However, the MPCC is impervious to such extra zeroes. The mean pixel intensity over the region of interest does not participate in the calculation of MPCC. The empty regions of the image outside the cell boundary cause corresponding zeroes in the 2D projected reference matrix. They affect neither the normalization condition (Eq. 4 in Methods) nor the calculated MPCC (Eq. 3). In the MPCC procedure, one need not worry about empty regions of the image matrices that lie outside the projected cell boundaries.

The caveats outlined here apply to essentially all types of cells or organelles. In principle, the central concept of MPCC can be generalized to other cell geometries, including irregular eukaryotic shapes. However, the MPCC will probably find its greatest use in bacterial cells, whose shapes are often quite uniform for given growth conditions. It is relatively straightforward to simulate appropriate 2D reference distributions for rod-shaped bacteria like *E*. *coli* and *B*. *subtilis*, using a spherocylinder as the simplified model. The problem becomes more difficult for other shapes, such as the spiral shaped *H*. *pylori*.

One possible purely experimental solution would be to co-image a large population of freely diffusing fluorophores that presumably map out the 2D projection of a 3D random distribution in the cell volume of interest. To test this concept on *E*. *coli*, we have imaged Kaede under the same growth conditions used to image the RNAP and HU spatial distributions of Fig. 4. Kaede is a non-native tetrameric fluorescent protein that diffuses freely in *E*. *coli* and appears to fill the cytoplasm uniformly (25). We imaged Kaede in 15 cells of length 3.6 to 3.8 μm, the same length bin used for RNAP and HU. The composite distribution of 54,719 spatial localizations from 8 of the 15 cells was pixelated to give an experimental estimate of **U͠^R^**. An estimate of **U͠^G^** was generated from the pixelated 2D projection of 66,301 Kaede copies from the other 7 cells. Using these experimentally generated matrices **U͠^R^** and **U͠^G^**, we calculated MPCC for the same RNAP and HU spatial distributions to be 0.56, 0.42 and 0.32 for chosen pixel sizes of 200 nm, 105 nm and 50 nm respectively. These completely experimentally derived MPCC values are similar to the MPCC values of 0.46, 0.39 and 0.25 obtained from simulation of **U͠^R^** and **U͠^G^** for the same respective pixel sizes. For cases in which it is difficult to simulate the 3D cell geometry, the experimental approach to generation of the reference matrices may prove useful.

## METHODS

As a first step towards generation of the matrices **U͠^R^** and **U͠^G^** in Eq. 3, a large number of molecules (here 100,000) are randomly distributed in a spherocylinder whose dimensions match those of the cells being imaged. We want **U͠^R^** and **U͠^G^** to have high signal-to-noise in each pixel. For a cell of length 3.5 μm and width of 0.82 μm, the choice of 200 nm for the pixel size results in 75 pixels in the cell. 100,000 molecules makes the mean occupancy 1333 molecules/pixel. An appropriate localization error σ is applied to each particle location in both *x* and *y* coordinates by sampling a Gaussian distribution with standard deviation σ, yielding the model “measured” location of each molecule, which is binned appropriately. For generating 3D random distribution of molecules corresponding to the red and green channels, the localization error applied is the same as that measured upon imaging fluorescent molecules in red (σ_*R*_) and green channels (σ_*G*_) respectively. The 2D projections along the *z* axis of these two 3D random distributions give the matrices **U^R^** and **U^G^**. The elements
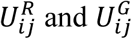
are positive integers.

Next the counts in individual pixels of **U^R^** and **U^G^** are normalized so that the total number of red and green molecules is equal to *N_R_* and *N_G_*, the total number of molecules imaged in each channel. This yields the normalized matrix **U͠^R^**:

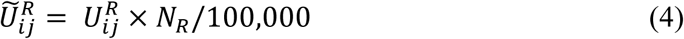

so that
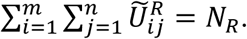
Similarly, **U^G^** is normalized so that the sum of all elements of **U͠^G^** is *N_G_*.

We then subtracted **U͠^R^** and **U͠^G^** from the corresponding image matrix under analysis, **R** and **G** respectively to obtain the unnormalized difference matrices **Δ^R^** and **Δ^G^**. We normalized **Δ^R^** and **Δ^G^** so that the sum of the squares of individual pixel values in the difference matrix is 1:

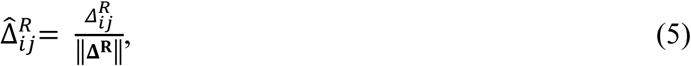

where
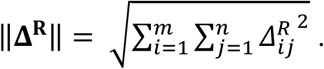
The resultant normalized 2D difference matrix **Δ̂^R^** has **∥Δ̂^R^∥** =1. The difference matrix **Δ^G^** in the green channel is similarly normalized to obtain **Δ̂^G^** such that **∥Δ̂^G^∥** = 1.

MPCC is obtained by taking the Frobenius inner product of the two normalized matrices **Δ̂^R^** and **Δ̂^G^**:

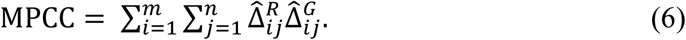

## Acknowledgments

This work was supported by a grant from the National Science Foundation (MCB-1512946 to JCW). We would like to acknowledge Mr. Sayar Karmakar for his insightful discussions on Pearson Correlation Coefficient.

## REFERENCES

1. Mondal J, Bratton BP, Li Y, Yethiraj A, Weisshaar James C. Entropy-Based Mechanism of Ribosome-Nucleoid Segregation in *E. coli* Cells. Biophys. J. 2011;100(11):2605–13.

2. Neeli-Venkata R, Martikainen A, Gupta A, Gonçalves N, Fonseca J, Ribeiro AS. Robustness of the Process of Nucleoid Exclusion of Protein Aggregates in *Escherichia coli*. J Bacteriol. 2016;198(6):898–906.

3. Manders EM, Stap J, Brakenhoff GJ, van Driel R, Aten JA. Dynamics of three-dimensional replication patterns during the S-phase, analysed by double labelling of DNA and confocal microscopy. J Cell Sci. 1992;103(3):857.

4. Pearson K. Mathematical contributions to the theory of evolution III. Regression, heredity, and panmixia. Philos Trans R Soc Lond B Biol Sci. 1896;187:253.

5. Schneider CA, Rasband WS, Eliceiri KW. NIH Image to ImageJ: 25 years of image analysis. Nat Methods. 2012;9(7):671–5.

6. Bolte S, CordeliÈRes FP. A guided tour into subcellular colocalization analysis in light microscopy. J Microsc. 2006;224(3):213–32.

7. Karunatilaka KS, Cameron EA, Martens EC, Koropatkin NM, Biteen JS. Superresolution Imaging Captures Carbohydrate Utilization Dynamics in Human Gut Symbionts. MBio. 2014;5(6):e02172–14.

8. Männik J, Wu F, Hol FJH, Bisicchia P, Sherratt DJ, Keymer JE, et al. Robustness and accuracy of cell division in *Escherichia coli* in diverse cell shapes. Proc Natl Acad Sci U S A. 2012;109(18):6957–62.

9. Strahl H, Bürmann F, Hamoen LW. The actin homologue MreB organizes the bacterial cell membrane. Nat Commun. 2014;5:3442.

10. Costes SV, Daelemans D, Cho EH, Dobbin Z, Pavlakis G, Lockett S. Automatic and Quantitative Measurement of Protein-Protein Colocalization in Live Cells. Biophys. J. 2004;86(6):3993–4003.

11. Dedecker P, Mo GCH, Dertinger T, Zhang J. Widely accessible method for superresolution fluorescence imaging of living systems. Proc Natl Acad Sci U S A. 2012;109(27):10909–14.

12. Dunn KW, Kamocka MM, McDonald JH. A practical guide to evaluating colocalization in biological microscopy. Am J Physiol Cell Physiol. 2011; 300(4):C723–C742

13. Earle Kristen A, Billings G, Sigal M, Lichtman Joshua S, Hansson Gunnar C, Elias Joshua E, et al. Quantitative Imaging of Gut Microbiota Spatial Organization. Cell Host & Microbe. 2015;18(4):478–88.

14. Skočaj M, Resnik N, Grundner M, Ota K, Rojko N, Hodnik V, et al. Tracking Cholesterol/Sphingomyelin-Rich Membrane Domains with the Ostreolysin A-mCherry Protein. PLOS ONE. 2014;9(3):e92783.

15. Wu Z, Tang M, Tian T, Wu J, Deng Y, Dong X, et al. A specific probe for two-photon fluorescence lysosomal imaging. Talanta. 2011;87:216–21.

16. George TC, Fanning SL, Fitzgeral-Bocarsly P, Medeiros RB, Highfill S, Shimizu Y, et al. Quantitative measurement of nuclear translocation events using similarity analysis of multispectral cellular images obtained in flow. J Immunol Methods. 2006;311(1):117–29.

17. Cabrera JE, Jin DJ. The distribution of RNA polymerase in *Escherichia coli* is dynamic and sensitive to environmental cues. Mol. Microbiol. 2003;50(5):1493–505.

18. Castaing B, Zelwer C, Laval J, Boiteux S. HU Protein of *Escherichia coli* Binds Specifically to DNA That Contains Single-strand Breaks or Gaps. J Biol Chem. 1995;270(17):10291–6.

19. Wang W, Li G-W, Chen C, Xie XS, Zhuang X. Chromosome Organization by a Nucleoid-Associated Protein in Live Bacteria. Science. 2011;333(6048):1445.

20. Nielsen HJ, Ottesen JR, Youngren B, Austin SJ, Hansen FG. The *Escherichia coli* chromosome is organized with the left and right chromosome arms in separate cell halves. Mol. Microbiol. 2006;62(2):331–8.

21. Biteen JS, Thompson MA, Tselentis NK, Bowman GR, Shapiro L, Moerner WE. Super-resolution imaging in live *Caulobacter crescentus* cells using photoswitchable EYFP. Nat Methods. 2008;5(11):947–9.

22. Subach FV, Patterson GH, Manley S, Gillette JM, Lippincott-Schwartz J, Verkhusha VV. Photoactivatable mCherry for high-resolution two-color fluorescence microscopy. Nat Methods. 2009;6(2):153–9.

23. Ando R, Hama H, Yamamoto-Hino M, Mizuno H, Miyawaki A. An optical marker based on the UV-induced green-to-red photoconversion of a fluorescent protein. Proc Natl Acad Sci U S A. 2002;99(20):12651–6.

24. Hayashi I, Mizuno H, Tong KI, Furuta T, Tanaka F, Yoshimura M, et al. Crystallographic Evidence for Water-assisted Photo-induced Peptide Cleavage in the Stony Coral Fluorescent Protein Kaede. J Mol Biol. 2007;372(4):918–26.

25. Bakshi S, Bratton Benjamin P, Weisshaar James C. Subdiffraction-Limit Study of Kaede Diffusion and Spatial Distribution in Live *Escherichia coli*. Biophys. J. 2011;101(10):2535–44.

26. Adler J, Parmryd I. Quantifying colocalization by correlation: The Pearson correlation coefficient is superior to the Mander’s overlap coefficient. Cytometry Part A. 2010;77A(8):733–42.

27. McDonald JH, Dunn KW. Statistical tests for measures of colocalization in biological microscopy. J Microsc. 2013;252(3):295–302.

28. Lagache T, Sauvonnet N, Danglot L, Olivo-Marin J-C. Statistical analysis of molecule colocalization in bioimaging. Cytometry Part A. 2015;87(6):568–79.

29. Owen DM, Rentero C, Rossy J, Magenau A, Williamson D, Rodriguez M, et al. PALM imaging and cluster analysis of protein heterogeneity at the cell surface. Journal of Biophotonics. 2010;3(7):446–54.

30. Perry GLW. SpPack: spatial point pattern analysis in Excel using Visual Basic for Applications (VBA). Environmental Modelling & Software. 2004;19(6):559–69.

31. Ripley BD. Tests of ‘Randomness’ for Spatial Point Patterns. Journal of the Royal Statistical Society Series B (Methodological). 1979;41(3):368–74.

32. Sengupta P, Jovanovic-Talisman T, Skoko D, Renz M, Veatch SL, Lippincott-Schwartz J. Probing protein heterogeneity in the plasma membrane using PALM and pair correlation analysis. Nat Methods. 2011;8:969.

33. Veatch SL, Machta BB, Shelby SA, Chiang EN, Holowka DA, Baird BA. Correlation Functions Quantify Super-Resolution Images and Estimate Apparent Clustering Due to Over-Counting. PLOS ONE. 2012;7(2):e31457.

34. Lee Steven F, Thompson Michael A, Schwartz MA, Shapiro L, Moerner WE. Super-Resolution Imaging of the Nucleoid-Associated Protein HU in *Caulobacter crescentus*. Biophys. J. 2011;100(7):L31–L3.

35. Fiolka R, Shao L, Rego EH, Davidson MW, Gustafsson MGL. Time-lapse two-color 3D imaging of live cells with doubled resolution using structured illumination. Proc Natl Acad Sci U S A. 2012;109(14):5311–5.

36. Jones SA, Shim S-H, He J, Zhuang X. Fast, three-dimensional super-resolution imaging of live cells. Nat Meth. 2011;8(6):499–505.

